# MenDEL: PCR Primer Design as Constrained Optimization Process

**DOI:** 10.1101/2022.06.26.496474

**Authors:** Sergei German, Leslie A. Mitchell, Antonio Vela Gartner, David Fenyö, Jef D. Boeke

**Affiliations:** Institute for Systems Genetics, NYU Grossman School of Medicine, New York, NY 10016, USA; Department of Biochemistry and Molecular Pharmacology, NYU Grossman School of Medicine, New York, NY 10016, USA; Department of Biomedical Engineering, NYU Tandon School of Engineering, Brooklyn NY 11201

## Abstract

**Motivation:** The synthesis of large DNA assemblies has applications in biotechnology, and can help us better understand genome biology. These large DNA assemblies are often constructed from many smaller DNA segments, and it is critical to assess that they are correctly assembled. One low cost and rapid method to ensure that the connection between each segment is correct is to use PCR with primer pairs that span assembly junctions. However, the design of PCR primers for large assemblies consisting of multiple segments, and therefore containing multiple assembly junctions, is a challenging process. Rule-based automation of the process often results in finding primers that satisfy general criteria, but are not necessarily the best fit for every particular junction.

**Results:** We have developed MenDEL – a web-based DNA design application, that provides a primer pair computation tool for multiple assembly junctions in such a way that for each junction we automatically pick the optimal pair of primers based on user specified constraints.

**Availability and Implementation:** The MenDEL application is available at https://mendel-isg.nyumc.org to registered users, and the code base for computing junction primers is available at https://github.com/MendelProject/PrimerOptimization.

## 1. Introduction

We describe here a module of an in-house DNA Design tool and LIMS called MenDEL, which stands for Mentored Design Environment and LIMS. One of the main purposes of MenDEL is to facilitate the assembly of large DNAs and a crucial part of that process is evaluating whether a given DNA assembly is “correctly assembled”, be it *in vitro* or *in vivo*. A typical assembly reaction yields many hundreds of clones or colonies that must be screened rapidly and comprehensively to identify putative “winner” carrying full-length assemblies. One simple and inexpensive approach to screen such colonies is to use polymerase chain reaction (PCR). Here, each assembly junction requires a pair of PCR primers to be designed and it is this process, to design not just one set of primers but a consistently performing set of primers for all the junctions, that is solved by the current approach. In theory, hundreds of primer pairs could be chosen that would span each junction of a given assembly, but in practice, identifying a set of primer pairs expected to function well under a single set of conditions and produce amplicons of roughly the same length is the goal.

Junction primers are oligonucleotides, usually 18-24 base pairs long. Primers are placed as inward-facing pairs on each side of an assembly junctions in order to ensure correct amplification (1). Fig.1 illustrates a short assembly with its segments. A pair of junction primers needs to be placed on both sides of each segment overlap and importantly, outside the actual overlap region itself. Success of polymerase chain reaction (PCR) to a large degree depends on the quality of these junction primers. Here, by quality of a primer we mean the ability of the primer to satisfy a number of user defined constraints – annealing temperature, length of the primer, distance from segment junctions, GC content, etc. There are a number of software applications that allow researchers to design individual primer pairs: Primer3 (2), Primer Plus (2), Primer BLAST (3). Another important primer design constraint for the application of DNA assembly evaluation is the uniqueness of the primer along the length of the target sequence.

**Figure 1.**
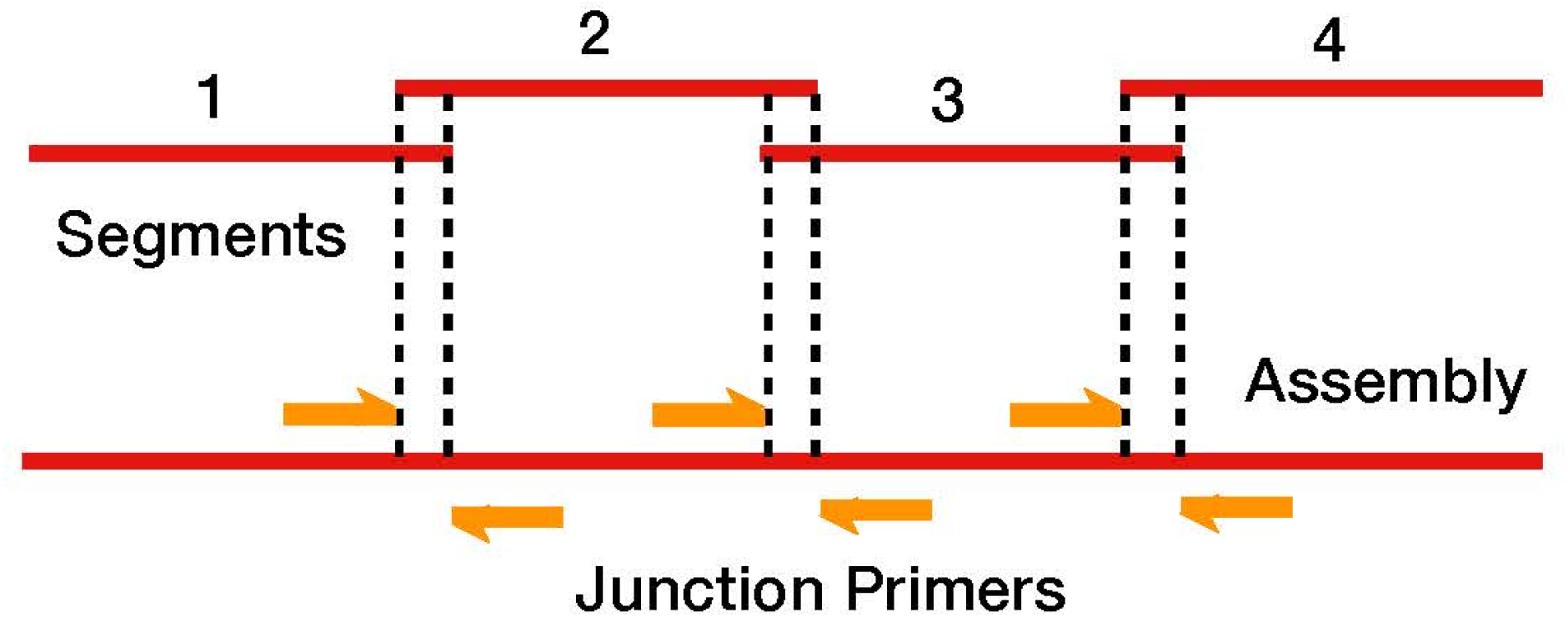
To test whether the assembly (bottom red line) is correct, pairs of PCR junction primers (orange arrows) are placed on both sides of each segment (top red lines) overlap (dashed lines).

Building DNA assemblies longer than a few thousand base pairs usually require joining together a large number of DNA segments by a variety of methods (4). Even though pairs of PCR primers for each segment junction could be designed individually, for example using the tools mentioned above, this is both time consuming and error prone. Another approach is to automate the process using a predefined set of rules, such as annealing temperature, amplicon length, GC content, etc. This automated constraint satisfaction approach solves some of the problems related to individual design of primer pairs – it is fast and less error prone. However, it introduces more subtle problems that may result in choosing less than optimal primer pairs for at least some of the junctions. As we set our constraints, we must make them relaxed enough to ensure that every junction has at least one pair of primers that satisfies these constraints. As a result of this constraint satisfaction approach, there might be junctions with multiple pairs of PCR primers that satisfy those constraints. Since the constraint satisfaction approach does not provide for ways to choose the “best” pair of primers, we might end up with primers that satisfy a relaxed general set of constraints but at the same time are not necessarily the best primers for each given junction.

To solve this problem, we propose a constrained optimization approach for automated primer design.

## 2. Implementation

### 2.1. Using a scoring scheme to find optimal primers

Constrained optimization (5,6) is an approach to finding minimal (or maximum) value of

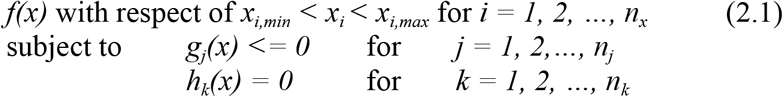

When applying this method to PCR primer selection we recognize that some of the constraints could be more important than the others, and, further, the relative importance of satisfying a particular constraint could change from one DNA assembly to the next. In addition, we must formally distinguish between logical and numeric (or range) constraints. An example of logical constraint would be an absence of a “hair pin” within a primer sequence. If there is no “hair pin”, then the constraint is satisfied and otherwise it is violated. An example of a range constraint would be the primer length - to satisfy this constraint, the primer length must be within specified range.

To find optimal primers for a given junction, we propose a “score” system – a primer with the highest score is chosen as an optimal primer. To compute this score we use the following equation:

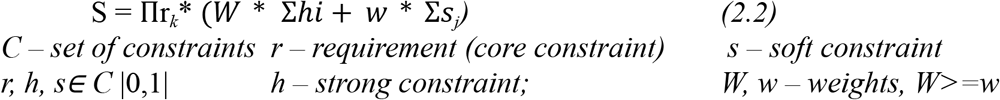

All primers must satisfy core constraints (requirement), those that satisfy additional constraints receive a higher score. For each junction the optimal primer - the one with the highest score - is chosen.

### 2.2. The MenDEL Primer Optimizer Package

A concrete implementation of primer design through applying constraint optimization is done in MenDEL’s optimizer package (the class diagram of this package is shown in Fig. 2).

**Figure 2.**
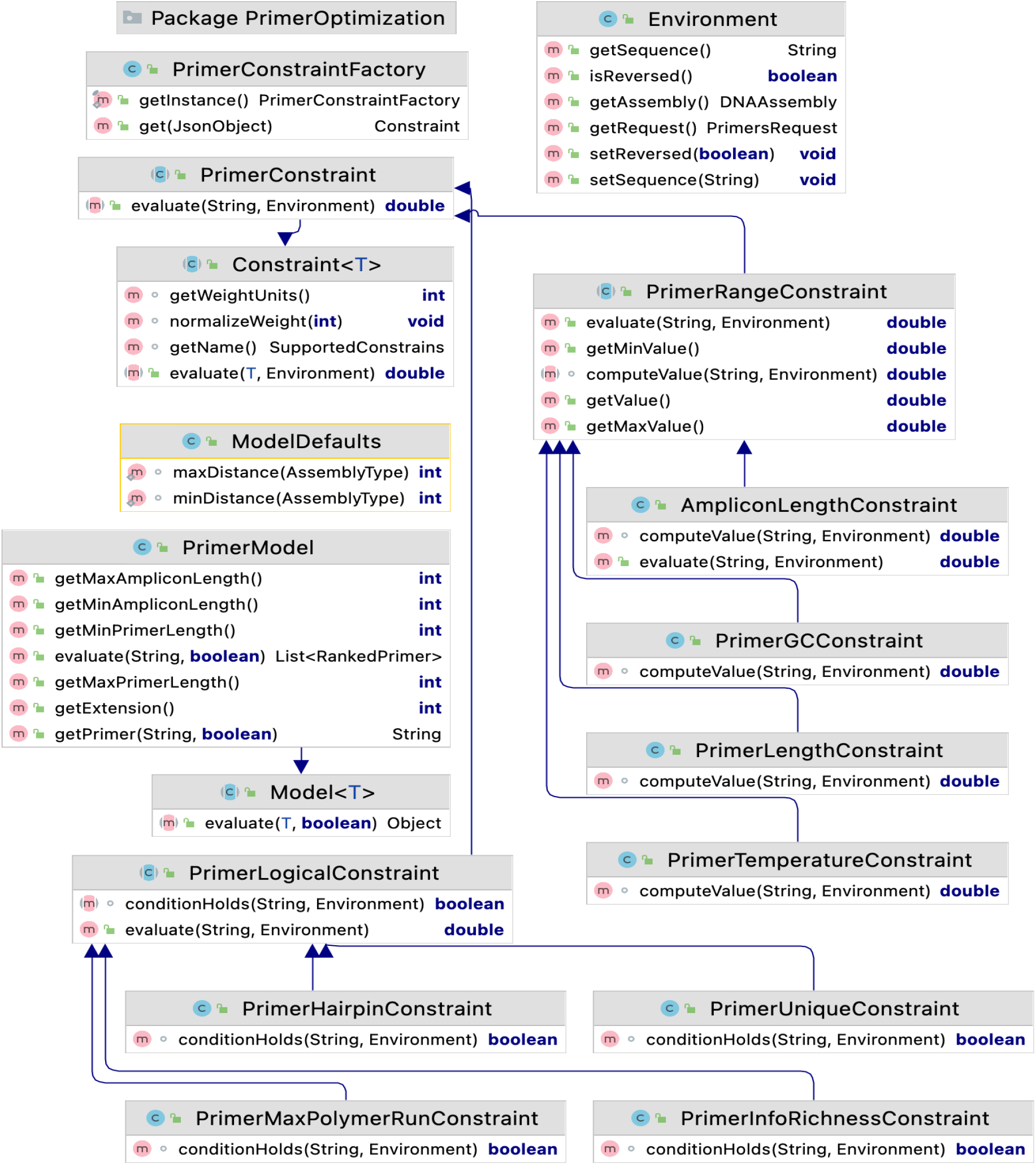
The MenDEL primer optimizer package class diagram.

The package implements a number of range constraints:

- PrimerGCConstraint - specifies that GC content of the primer is in the specified range
- AmpliconLengthConstraint - checks whether the amplicon (length of the sequence between forward and reverse primers) is in the specified range
- PrimerLengthConstraint - Primer length is in specified range
- PrimerTemperatureConstraint - specifies the desired annealing temperature range

Logical constraints are implemented in these classes:

- PrimerHairpinConstraint - primer does not form a ‘hairpin’ secondary structure that will inhibit success in PCR
- PrimerUniqueConstraint - primer is unique within the assembly for PCR (first 15 and last 15 base pairs, as well as their reverse complements are unique)
- PrimerMaxPolymerRunConstraint - primer does not contain a polymer run (i.e. no more than 5 A, C,G or T in a row)
- PrimerInfoRichnessConstraint - specifies that sequence of each primer have a minimum 10% representation of each of the bases

Computational model – scoring system - is implemented in PrimerModel class. Environment class represents specific DNA assembly, DNA segment sequences, etc. for which primers are computed. Package class hierarchy allows users to expand the constraint universe by providing new classes that implement PrimerRangeConstraint or PrimerLogicalConstraint and register them with PrimerConstraintFactory.

### 2.3. MenDEL – Junction Primer Design module

The junction primer design module, based on the constraint optimization model, is integrated into the MenDEL application. The current implementation (see **Fig. 3**) takes advantage of a drag-and-drop user interface. By moving constraints from the Constraint Palette to one of the buckets – Requirement, Strong Preference and Soft Preference - the user can effectively build the model equation 2.2. By specifying assembly attributes (Study, Project, Assembly name) users establish the computational environment. By applying the model and environment, MenDEL’s engine then computes optimal primer pairs for each of the assembly’s junctions.

**Figure 3.**
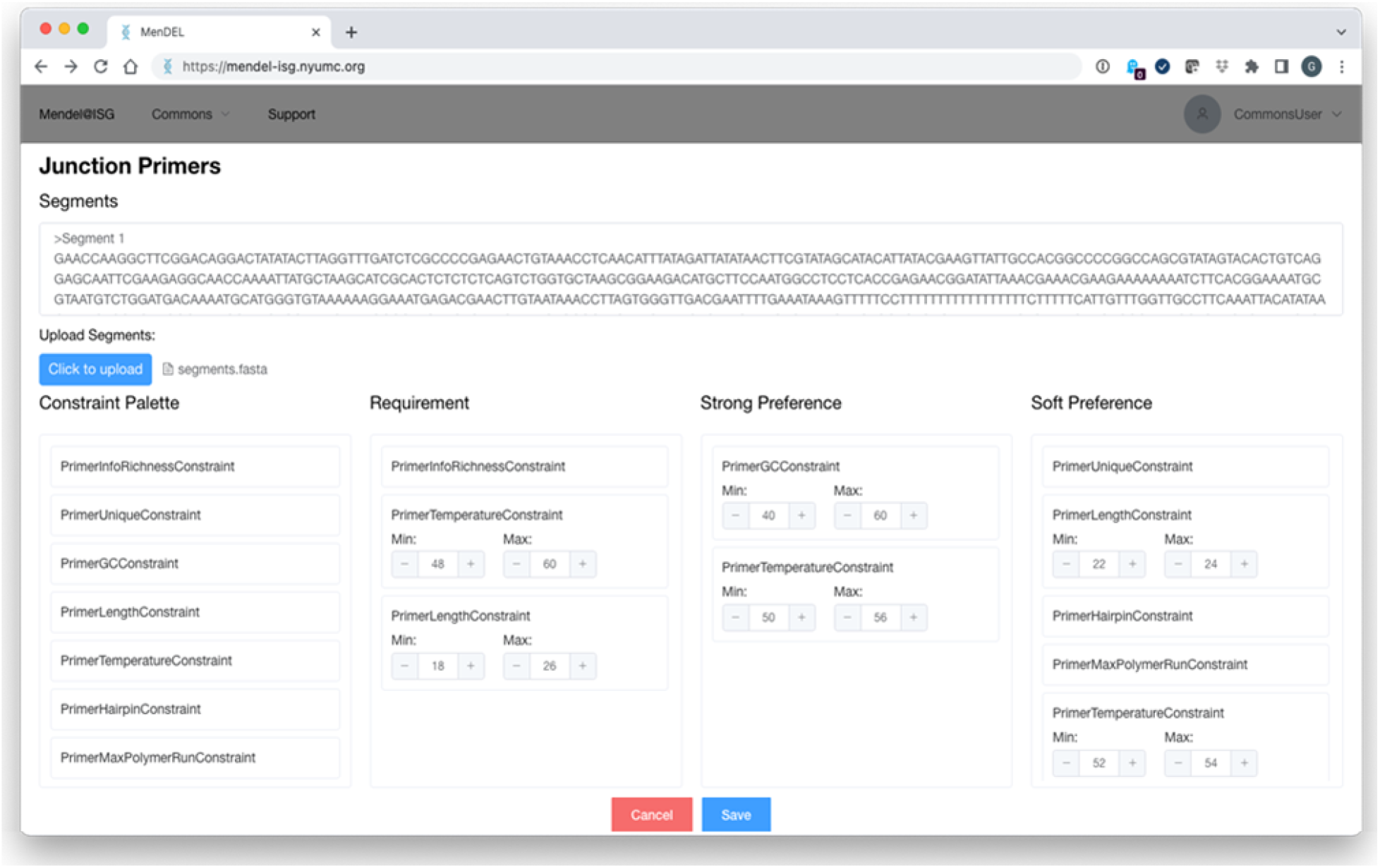
User interface for the optimized junction primers module.

By uploading the segments and distributing constraints from the Palette to desired buckets (Requirement, Strong and Softy Preferences), a user effectively provides a computational environment and model for applying optimized constraints approach for Junction Primer Design.

We have shown (Figure 4) that PCR primers designed using default parameters (depicted on Figure 3) correctly amplify xx/yy junction fragments using the GoTaq Green system on a 95 kb BAC clone template.

**Figure 4.**
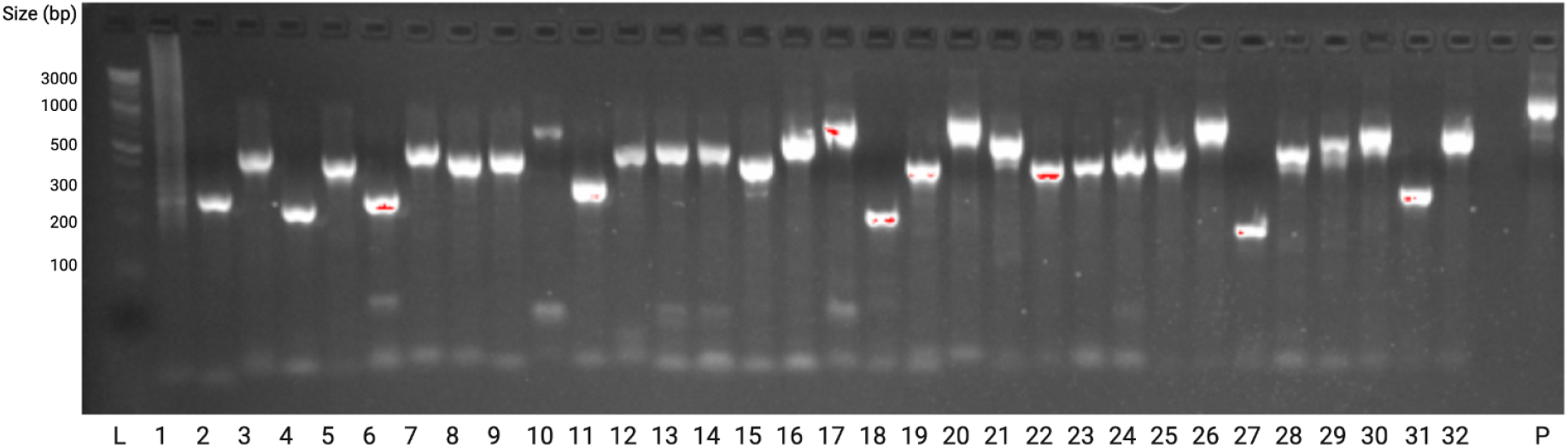
To assess MenDEL’s junction primer module we simulated the segmentation of a previously built 96 kb DNA assembly and designed 32 pairs of junction primers to be used in parallel PCR reactions with identical conditions. 5 ng template were used in each reaction. L – 1 kb Plus DNA Ladder, 1-32 – PCR with primer pairs for junctions of the assembly, P – positive control primer pair.

## 3. Conclusion

The constraint optimization approach to junction primer design allows finding optimal – in respect to the highest score – primers for each of an assembly’s junctions. It combines a high level of flexibility of primer design, associated with manual design of each primer, with low rate of human errors and high efficiency of computation, associated with automated primer design. These characteristics make constraint optimization the approach especially valuable for assemblies with a large number of junctions.

Implementation of this approach in the MenDEL application provides an architecture that allows ease of support and enhancement – users and developers can easily add new constraint classes, provide their own implementation for existing constraints or computational models. The drag-and-drop user interface allows both experienced and novice users to easily perform the optimization and fine-tune it according to their preferences.

Efficiency of computation, flexibility of primer design and of finding the optimal primers for each junction, ease of use, and possibility of customization makes constraint optimization a promising approach for PCR primer design.

## Acknowledgments

This work was supported in part by NIH/NHGRI grant 1RM1HG009491.

